# Heterosis from local drift load is likely insufficient to favor reversions to outcrossing

**DOI:** 10.1101/273946

**Authors:** Alexander Harkness, Emma E. Goldberg, Yaniv J Brandvain

## Abstract

The evolutionary trajectory from cross-to self-fertilization is widely documented in nature, but results from several taxa also suggest that outcrossing may evolve in a formerly selfing population. Population genetic theory explains that selfing can evolve when its advantages overcome its immediate cost of inbreeding depression, but that this process will not run in reverse because a self-fertilizing population purges itself of inbreeding depression. That is, the primary short-term advantage of cross-fertilization over self-fertilization depends on the existence of deleterious alleles exposed upon inbreeding. Here, we explore whether outcrossing can evolve in selfing populations if allelic variation exists as divergence among populations. We consider two monomorphic populations of entirely self-fertilizing individuals, introduce a modifier allele that increases the rate of cross-fertilization, and investigate whether the heterosis among populations is sufficient for the modifier to invade and fix. We find that, despite an initial increase in the frequency of the outcrossing modifier, its fixation is possible only when populations harbor extremely large unique fixed genetic loads. These rare reversions to outcrossing become more likely as the load becomes more polygenic, or when the modifier appears on a rare background, such as by dispersal of an outcrossing genotype into a selfing population.

## Introduction

The transition from a predominantly outcrossing to a predominantly self-fertilizing (selfing) mating system is a very common evolutionary event among angiosperms, supported, for example, by many observations of predominantly selfing taxa nested within outcrossing taxa or retaining outcrossing adaptations (Stebbins 1974). A large body of population genetic theory describes conditions under which this common transition is expected to occur (Fisher 1941; Kimura 1959; Nagylaki 1976; Lloyd 1979; Lande and Schemske 1985). The reverse transition, from selfing to outcrossing, is substantially less common, but its occurrence has been hypothesized in a some taxa (Barrett and Shore 1987; Olmstead 1990; Bena et al. 1998). However, the evolution of outcrossing in a formerly selfing population has received relatively little theoretical attention and it is therefore unclear when, if ever, it could occur.

The theoretical neglect of possible reversion from selfing to outcrossing is likely due to the absence of an obvious disadvantage of selfing in historically selfing populations. Selfing exposes recessive alleles underlying inbreeding depression to removal by selection (“purging” of inbreeding depression), which enables evolution of greater selfing and removal of more inbreeding depression (Lande and Schemske 1985). Furthermore, new recessive deleterious mutations would be rapidly exposed to selection in a highly selfing population, and inbreeding depression would therefore be unlikely to re-accumulate. With no short-term disadvantage, high levels of selfing are expected to persist indefinitely within a population.

Still, several authors (Barrett and Shore 1987; Olmstead 1990; Bena et al. 1998) have claimed examples of reversion from predominant selfing to outcrossing—how, then, could these hypothesized reversions have occurred? The first possibility is for inbreeding depression to persist or re-accumulate in a predominantly selfing population. Some inbreeding depression can indeed survive the purging process (Willis 1999). Intermediately selfing populations in nature do harbor inbreeding depression equivalent to that of predominantly outcrossing populations on average, but predominantly selfing populations seem to have lost most of their inbreeding depression through purging or fixation of deleterious alleles (Winn et al. 2011). The second possibility is for an alternative disadvantage of selfing to exist despite the depletion of inbreeding depression. One cause would be if some deleterious alleles are stochastically fixed by drift rather than being purged by selection. If these deleterious alleles are partially or fully recessive, outcrossing but not selfing individuals can mask them by hybridizing with another population. Here we present and evaluate the hypothesis that this “local drift load”—the genetic load of fixed deleterious alleles peculiar to a local population (Whitlock et al. 2000)—is sufficient to favor outcrossing in otherwise selfing populations.

Local drift load is particularly likely to accumulate in highly selfing populations. Although selfing exposes recessive alleles to selection, it relaxes selection overall. Relaxed selection occurs partly because non-independent gamete sampling reduces effective population size, and especially because reduced effective recombination allows more hitchhiking by deleterious alleles (Barrett et al. 2014). Drift load is local because different alleles have fixed by chance in each population. Outcrossing individuals could potentially avoid their local drift load by crossing with members of populations lacking the fixed deleterious alleles present in their own population, whereas selfers could not. Assuming that the fixed deleterious alleles are recessive on average, heterozygous hybrids will mask their fixed load and will show heterosis in fitness. Any differences in deleterious allele frequency can contribute to heterosis (Whitlock et al. 2000), but we focus on the conceptually simple extreme case of fixation versus absence.

Heterosis is any increase in a trait value for hybrids above some expectation based on the parents. Although there are multiple ways to quantify heterosis, we follow Busch (2006) in defining heterosis as the relative excess of the expected fitness of offspring produced by among-population crosses over that of offspring produced by within-population crosses. Because heterosis is only attainable through outcrossing, local drift load may favor alleles that increase the outcrossing rate. If predominantly selfing populations are more susceptible to local drift load, they should also show more heterosis than predominantly outcrossing populations. Empirical heterosis estimates in populations of different census sizes (Heschel and Paige 1995), levels of isolation (Richards 2000), and breeding systems (Busch 2006) generally support the proposition that isolated, effectively small populations accumulate greater heterosis than effectively large, connected populations (but see Ouborg and van Treuren 1994). These results support the expectation that inbreeding and selfing lead to local drift load. It is therefore realistic to expect that individuals in some predominantly selfing populations would benefit from hybridization.

To investigate the conditions under which greater outcrossing can evolve from predominant selfing, we simulate a population genetic model of the invasion of a modifier mutation that increases the rate of outcrossing in a pair of completely selfing populations that have recently come into secondary contact. In a meta-population model, Theodorou and Couvet (2002) previously showed that the advantage of outcrossing increases with population isolation partly because isolation allows the accumulation of local drift load and heterosis. Our model asks if this process can be extended to the secondary evolution of outcrossing in two formerly isolated selfing populations which have each independently accumulated their own local drift load. Our model is thus suited to evaluating whether secondary contact can extricate a population from what within-population evolution would suggest is an absorbing state.

Our results verify that local drift load initially favors outcrossing, but they also show that in most cases this advantage is quickly lost. Therefore, while the frequency of an outcrossing modifier allele initially increases under a wide portion of parameter space, it is ultimately lost before it achieves fixation unless heterosis is unreasonably strong. To better understand when and how the local drift load can result in a reversion to outcrossing, we investigate how the polygenicity of this load and the relative sizes of the populations can make this reversion more or less probable.

## Model

To determine whether heterosis can favor outcrossing, we considered an idealized model that seems especially likely to do so. Thus our results represent a “best case scenario” for the secondary evolution of outcrossing. We begin with a conceptual outline and later describe the specific simulation steps. All simulations and numerical iterations were performed using custom R scripts (provided as Supplementary Information; R Core Team 2016).

### General model description

#### Population history

We began with two selfing populations with unique sets of fixed alleles that affect viability, either beneficial mutations fixed by selection or deleterious mutations fixed by chance in spite of selection. One population possessed the superior allele at half of the viability loci, while the other population possessed the superior allele at the other half. By requiring the ancestral genotypes to be of exactly equal fitness, we avoided competitive exclusion of one population by the other. Our scenario could correspond to a vicariance event, such as the creation of a physical barrier, that split a population in half. Selfing could have been inherited from this common ancestor, or it could have evolved in both populations after they split.

The two populations then came into secondary contact. We assumed that there was unrestricted migration and that the populations wer of equal size (though we relax the latter assumption later). Instead of explicitly modeling the amount of migration between two separate populations, we treated the populations after secondary contact as a single merged population composed of individuals from both parent populations. The *F*_1_ individuals were heterozygous at all loci at which their parents were fixed for divergent alleles. We assumed that all inferior alleles were completely recessive in order to maximize heterosis (Whitlock et al. 2000), and to reflect the fact that strongly deleterious alleles tend to be recessive in nature (Mackay et al. 1992; Caballero and Keightley 1994). Our model describes evolution from the time of secondary contact onwards.

#### Mating system

Only outcrossing can produce *F*_1_s. We considered a rare “outcrossing allele,” *M*, at a modifier locus unlinked to any loci under direct selection. This allele may have arisen after contact or shortly before. The common, wild-type “selfing allele” at this locus is denoted *m*. The outcrossing phenotype we modeled best corresponds to a morphological trait that prevents self-pollination, rather than to self-incompatibility, because it caused random rather than disassortative mating. Under our main model *M* was completely dominant, although we later examine an additive version. Carriers of *M* fertilized their ovules with pollen drawn at random from the pollen pool, whereas *mm* homozygotes self-fertilized all their ovules. In the terminology of Lloyd (1992) seed discounting is complete. Of the different modes of selfing Lloyd (1992) defined, ours was similar to prior selfing in that the fraction of ovules selfed was independent of the supplies of self and outcross pollen. However, each of Lloyd’s modes of selfing assumed a limited supply of ovules for each dam, while ours did not.

The modifier genotype only controlled mating, and it had no direct effect on viability or fecundity. Therefore, selfing in our model lacked the advantage of reproductive assurance (Lloyd 1992) and the disadvantage of pollen discounting (Nagylaki 1976; Holsinger et al. 1984; Lloyd 1992). Fisher (1941) showed under these assumptions that self-fertilization transmits 50% more gametes than outcrossing because the additional siring success on selfed ovules does not detract from outcross siring success.

### Invasion condition

We obtained analytically the conditions under which an outcrossing allele was expected to increase in frequency initially in an infinite population with two viability loci. Viability selection was chosen so that selection would occur after mating, and so the modifier’s effects on progeny genotypes would be exposed to selection in a single generation. Otherwise, the change in allele frequencies over a generation could not be used as a criterion for invasion because it would reflect the effect of transmission advantage but not of offspring fitness. The model is described more explicitly in the next section, and the Appendix contains the invasion condition (Inequality 2) and its derivation. Essentially, the selfing allele *m* always increases in frequency in the seed pool before selection because of its transmission advantage. For the outcrossing allele *M* to increase in frequency by the next generation, *M*-carrying offspring must have sufficiently greater average viability that *m* decreases in frequency among adults despite having increased in frequency among seeds. However, numerical iteration showed that, at least in the deterministic case, two loci were too few to allow *M* to reach fixation even for extremely large *s* (Fig. 1). In contrast, recombination would take longer to break up the repulsion-phase linkage disequilibrium among greater numbers of loci, which could delay the purging of the load. Are greater numbers of loci sufficient to favor outcrossing?

**Figure 1:**
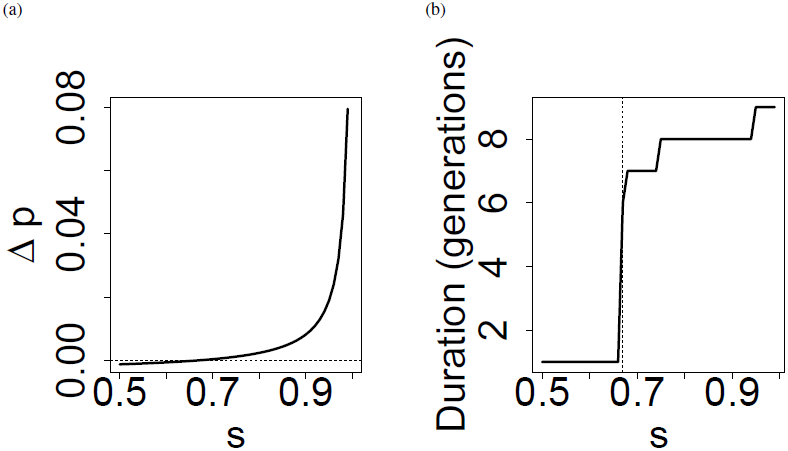
Invasion condition for the outcrossing modifier in the deterministic model. The population begins with equal frequencies of two viability genotypes, each fixed for a single inferior allele at a different locus. The outcrossing modifier, *M*, initially exists only in *Mm* heterozygotes, at frequency 1/200. (a) *M* invades in the first generation (Δ*p >* 0) only when selection is sufficiently strong (large *s* for each viability locus).Despite this initial increase in frequency, *M* is lost in subsequent generations (not shown). (b) The transit time during which *M* is segregating before its loss increases with the strength of viability selection. The dashed horizontal line corresponds to (Δ*p* = 0. The dashed vertical line shows the threshold *s* required for *M* to increase in frequency in the first generation, calculated from Inequality 2. Extinction is defined as a frequency less than 1*/*200, and fixation as frequency greater than 199*/*200, to correspond to the loss of the final copy of an allele in a diploid population of 100 (like in the stochastic model, described later).

### Simulation model

We next used a simulation model to investigate the dynamics and fixation probability of the outcrossing modifier, allowing for more viability loci. Although numerical iteration could track long-term behavior of deterministic recursions, it would only account for the expected outcome each generation. However, we were interested in possible transitions to greater outcrossing rates even if increased outcrossing is not a guaranteed consequence of heterosis. In particular, we wished to investigate whether increased outcrossing was substantially more probable with more heterosis. Additionally, deterministic iteration ignores the disadvantages of non-recombination (and thus of selfing) caused by stochasticity in the haplotypes that are produced and survive (Muller 1964; Hill and Robertson 1966), and these stochastic effects could play a major role in the evolution of outcrossing. We therefore estimated by simulation the probability that a new mutation that increases outcrossing would fix in a population that was previously completely selfing. We did this under multiple models of the magnitude and genetic architecture of heterosis. Specifically, we varied the selection coefficient *s* at each viability locus and the number of loci *L* at which each parental population was fixed for an inferior allele. We required each fixed deleterious allele in one parental population to be balanced out by a fixed deleterious allele at another locus in the other parental population in order to prevent one parental genotype from rapidly outcompeting the other.

We next describe the steps of the simulation model. (The analytic model used above to obtain the invasion condition is identical, except that it assumes infinite population size and only two, unlinked viability loci.) We assumed that the viability loci were evenly spaced on each of ten chromosomes, and that each chromosome carried the same number of viability loci. The exception is the first chromosome, which took the remainder if the number of loci was not divisible by the number of chromosomes. The relative positions of the superior alleles allotted to each genotype were randomized at the beginning of each simulation. We neglected mutation during the invasion process because we were only interested in heterosis built up before contact. For each model, we recorded the proportion of simulations resulting in the fixation of the modifier.

#### Mating

Each generation, a seed pool ten times the size of the adult population was generated. For each seed to be generated, a dam was chosen at random. If the dam had a selfing phenotype (genotype *mm*), it was also the sire. If the dam had an outcrossing phenotype (genotype *MM* or *Mm*), a random individual was chosen to be the sire. Recombination among loci and random sampling of one recombinant from each parent generated the parental gametes. The parental gametes were then united to create the diploid seed genotype. Although the number of outcrossed offspring that were hybrids was random, the probability of zero hybrids was small. A few failures to produce any hybrids were therefore not likely to dominate the results.

#### Recombination

We placed the viability loci on a genome of ten linear chromosomes. The outcrossing modifier locus was unlinked to any of the viability loci. In each region between viability loci, recombination occurred with probability *r* independently of recombination elsewhere in the genome. Viability loci were evenly spaced along each chromosome such that *r* was the same between any two neighboring loci. Loci on separate chromosomes assorted independently. The positions at which each population was fixed for the superior allele is randomized each simulation.

#### Selection

Soft viability selection among seeds generated the next adult generation. Such a scenario is similar to competition among seeds or seedlings in a small plot: although all individuals would be physically capable of survival on their own, competition eliminates the worse competitors. 10% of seeds were sampled without replacement to survive to adulthood. Seed *i*’s probability of being sampled was weighted by its viability 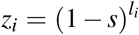, where *s* is the selection coefficient at each viability locus, and *l*_*i*_ is the number of viability loci homozygous for the inferior allele in individual *i*. Viability selection was chosen in order to match the deterministic model.

#### Fixation proportions

We found that 1000 generations was sufficient to allow the outcrossing allele to run its course to fixation or extinction in preliminary simulations. However, two parameter combinations (*L* = 50, *s* = 0.1, *r* = 0.5 and *L* = 14, *s* = 0.7, *r* = 0.1) still failed to complete, so they were re-run. To simulate a new mutation, we set the initial frequency of *M* to 1*/*2*N*, where *N* is the population size. We chose a small population size of ***N*** = 100 because the simulation scaled poorly with ***N***, but we compensated by performing enough replicates to observe the diversity of outcomes. We chose a number of trials per parameter combination sufficient that we would expect ten chance fixations of a unique neutral variant (2000 trials per parameter combination for an initial frequency of 1*/*2***N***). Fixations of *M* more or less frequent than the neutral expectation would imply positive or negative selection on *M*, respectively. The expected number of fixations of a hypothetical neutral allele was chosen to be greater than one so that variation below the expectation could still be detected, but ten was an arbitrary choice. We varied parameters from ***L*** = 5 to 50, and *s* = 0.1 to 0.9. These extremely large selection coefficients were expected to favor outcrossing by creating large magnitudes of heterosis. We considered two models of recombination frequency between neighboring viability loci: *r* = 0.5 and 0.1.

#### Transient dynamics

We ran additional simulations of the main model in which we tracked the frequency of *M* through time rather than only the final outcome. Based on the results of previous simulations, we chose parameter combinations with *L* = 5, 25, and 50 and *s* = 0.3 to cover a wide range of fixation probabilities. The allele frequency of ***M*** was recorded every generation, and the duration of the simulation and the population mean fitness were recorded at the end of each simulation.

#### Additional models

In our main model, we assumed that the two initial genotypes began at exactly equal frequencies, and that *M* was completely dominant. To investigate the effects of these assumptions, we simulated two additional scenarios. In the first, the two initial populations had unequal size. This might be analogous to cases where one population receives rare migrants from an allopatric population, for example. In this scenario, *s* = 0.1 for all parameter combinations. In the second, we assumed additive gene action at the modifier locus. Complete dominance at the viability loci was chosen to maximize heterosis, so we did not investigate other forms of gene action at these loci. In this scenario, *s* = 0.3 for all parameter combinations. In these variant models, we allowed free recombination (*r* = 0.5) and consequently simplified the genome structure to two chromosomes.

## Results

### Outcrossing mutation never fixed in a three-locus deterministic model

The threshold value of the selection coefficient, *s*, above which the outcrossing modifier, *M*, was expected to increase in the first generation was slightly below 0.67 according to Inequality 2 when the initial frequency of *M* is 0.005. The threshold approached 2*/*3 as the frequency of *M* approached zero. Numerical iteration for a single generation confirmed that *M* was lost in a single generation below the threshold and was lost after six or more generations above it (Fig. 1). Thus, the outcrossing modifier never fixed in the deterministic model. All subsequent results are for the stochastic simulation model.

### Outcrossing mutation could fix under restricted circumstances

A rare outcrossing mutation fixed in some of the simulations (Fig. 2). For some parameter combinations, it fixed more often than the expectation (0.005) for a neutral, initially unique allele. For much of parameter space, however, it fixed less often than the neutral expectation. In many parameter combinations, it never fixed at all. The outcrossing allele fixed when many loci were under selection and/or when there was strong selection at each locus.

**Figure 2:**
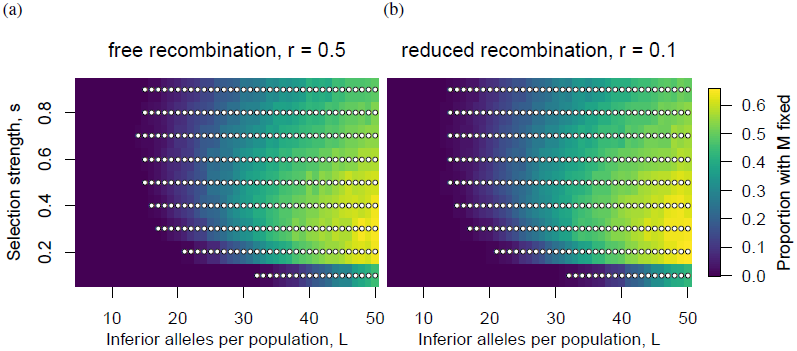
Outcrossing allele fixation proportions. Plotted is the proportion of simulations in which *M* fixed, for different combinations of number of viability loci and selection strength at each of those loci. Dots mark parameter combinations for which fixation was more frequent than the neutral expectation. Panels (a) and (b) differ in the amount of recombination between the viability loci. Fixation proportion increases with the number of loci under selection, at a rate that depends on the selection coefficient. Fixation increases most steeply with number of loci for intermediate selection coefficients. High proportions of fixation can be achieved at lower levels of heterosis for more polygenic loads of loci of smaller effect. Loose linkage has little effect.

### No effect of loose linkage

There was no qualitative difference between the results of simulations with free recombination between viability loci (*r* = 0.5) and those with reduced recombination (*r* = 0.1; Fig. 2). The outcrossing allele fixed more often than the neutral expectation in five more parameter combinations when *r* = 0.1 (303 parameter combinations) than when *r* = 0.5 (298 parameter combinations). The maximum difference in fixation proportion for any parameter was 0.06. The maximum fixation proportions were 0.6545 and 0.6455 for *r* = 0.1 and *r* = 0.5, respectively. Differences showed no consistent pattern with respect to *s* or *L*. All further results are reported for *r* = 0.5.

### Intermediate selection favored outcrossing

The fixation proportion was greater for larger than for the smallest values of the selection coefficient, *s* (Fig. 2). However, the relationship was not monotonic. Fixation proportion increased sharply going from *s* = 0.1 to *s* = 0.2. Fixation proportion reached a maximum somewhere in the range from *s* = 0.2 to 0.4, but the exact value depended on the number of viability loci. At *L* = 50, fixation was maximized at *s* = 0.3.

### Polygenic load favored outcrossing

The outcrossing allele fixed more often when the ancestral populations differed by many fixed viability loci. Holding the selection coefficient constant, the fixation proportion generally increased with the number of viability loci (Fig. 2). Fixation was maximized at the greatest number of viability loci modeled: *L* = 50. This effect was not due simply to heterosis being stronger with more loci, though. The outcrossing allele fixed more often under a polygenic heterosis than an oligogenic heterosis of equal magnitude (Fig. 3).

**Figure 3:**
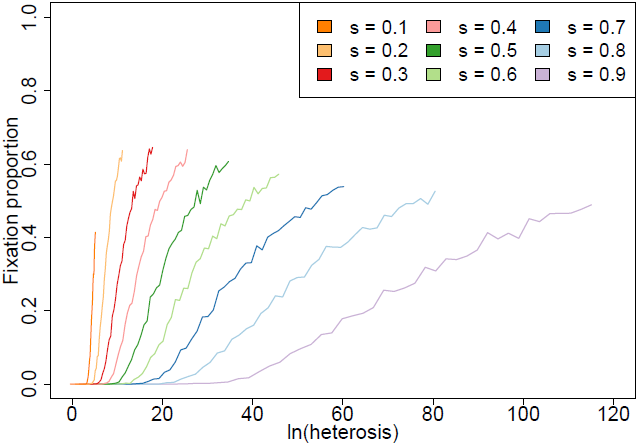
Effect of heterosis magnitude and architecture on fixation. Data are identical to those of Fig. 2a. Fixation generally increased with increasing heterosis. Comparing loads of equal magnitude, more polygenic loads (with smaller selection coefficients per locus) more often resulted in fixation than did more oligogenic loads.

### Outcrossing was sometimes lost after initially invading

The outcrossing modifier, *M*, often ultimately went extinct even when it initially increased in frequency (Fig. 4). There was a qualitative difference between parameter combinations in which *M* sometimes fixed and those in which it never fixed. In parameter combinations in which *M* never fixed, the initial increase in frequency was much smaller (Fig. 4, left column). Loss was rapid when it occurred, regardless of the frequency of *M* before the decrease in frequency began.

**Figure 4:**
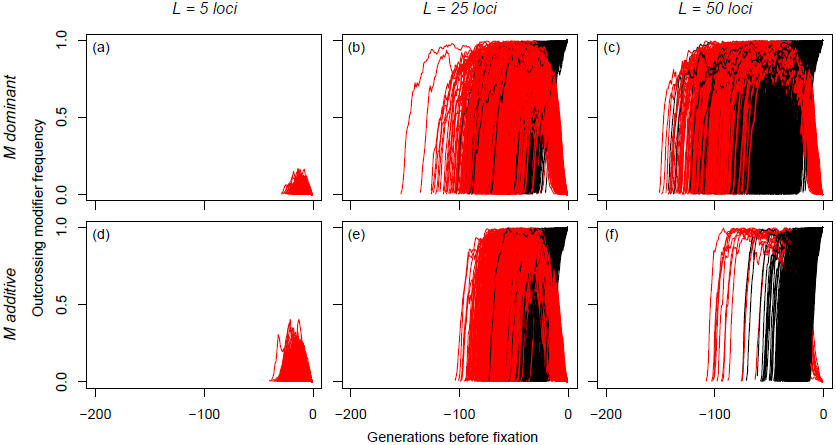
Allele frequency trajectories of the outcrossing modifier, *M*. Each trajectory represents a single simulation run. In each run, all individuals are initially homozygous for the selfing allele *m*, except that a single *M* mutation appears in a random individual in the first generation. Black trajectories resulted in fixation of *M*; red trajectories resulted in fixation of *m*. Time is measured relative to the final generation, called 0. Columns correspond to different numbers of viability loci. Rows correspond to different gene action of the modifier: dams that are heterozygous at the modifier locus outcross with probability 1 (top row) or 0.5 (bottom row). *s* = 0.3 for all parameter combinations.

### Additive modifiers fixed more often

Compared to dominant outcrossing alleles, alleles that additively increased the outcrossing rate did not slow their invasion at high frequencies before reaching fixation (Fig. 4). Such additive alleles spent less time segregating, and they more rarely experienced a sudden reversal in allele frequency trajectory. This resulted in an increased fixation probability for additive alleles. At *L* = 5, *M* never fixed. At *L* = 25, *M* fixed in 343 (dominant) or 598 (additive) out of 2000 trials. At *L* = 50, *M* fixed in 1292 (dominant) or 1599 (additive) out of 2000 trials.

### Ultimate mean fitness and time to extinction were bimodal

All simulation durations and final mean fitnesses are summarized in Fig. S1, Fig. S2, and Fig. S3. Simulations resulting in fixation of *M* most often ended with high but not maximal mean population fitness. Simulations resulting in loss of *M* showed a bimodal pattern with the largest number eventually ending with very high fitness and a smaller number rapidly ending with very low fitness (Fig. 5). The shortest and longest runs resulted in extinction of *M*, whereas runs resulting in fixation lasted an intermediate duration (Fig. 5 insets). Purging was more complete in the longer runs than in the shorter ones (Fig. S4): the mean final mean fitness of all simulations with *L* = 50 was 0.81, while the mean for the subset with below-average duration was 0.69. The modal duration was one generation for all six of the parameter combinations that were tracked through time (***s*** = 0.3, *L* = 5, 25, or 50, additive or dominant *M*). Only the results for the parameter combination with *L* = 25 and dominant *M* were plotted (Fig. 5), but this scenario was broadly representative. The exceptions were that at *L* = 5, no fixations were observed and the ultimate fitness had one mode concentrated near zero (Fig. S1), and that in the additive model with *L* = 50, most extinctions of *M* were rapid (Fig. S3).

**Figure 5:**
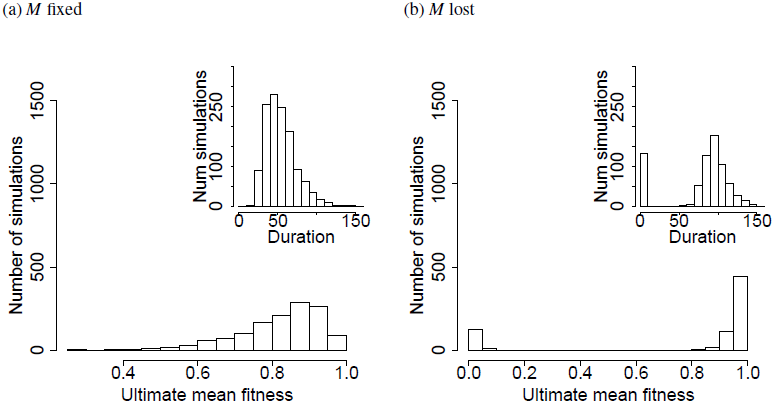
Distribution of ultimate mean fitnesses. The mean fitness of the population was recorded in the final generation of simulated invasions of *M*. The simulation runs are subsetted into those that resulted in fixation of *M* (a) or loss of *M* (b). The inset in each panel is the distribution of simulation durations conditional on the same outcome (fixation or loss). Data are identical to those in Fig. 4, with parameter values *L* = 50 and *s* = 0.3, and *M* completely dominant.

### Rare genetic background favored outcrossing mutation

In the simulations of unequal initial frequencies of the two ancestral genotypes, the outcrossing allele fixed more often when it arose on the rarer genetic background (Fig. 6). However, this effect was only visible when there were many viability loci (*L* = 50) because no fixations occurred at *L* = 5 or 25 when *s* = 0.1.

**Figure 6:**
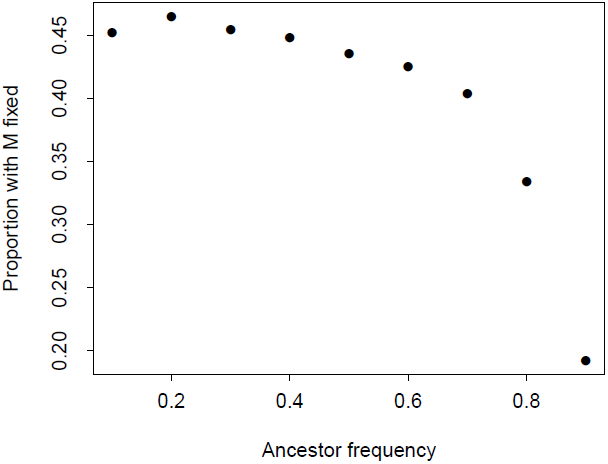
Fixation proportion with initially asymmetric viability genotype frequencies. Simulations were run in which the outcrossing modifier originated on a rarer or more common genetic background. These correspond to an ancestor frequency less than or greater than 0.5, respectively. Parameter values are *s* = 0.1, *r* = 0.5, and *L* = 50, and *M* is dominant. No fixations occurred for *L* = 5 or 25 (not shown).

### Overall interpretation

Some parameter combinations resulted in fixation proportions for *M* greater than the expected proportion of 1*/*2*N* for a unique neutral mutation, suggesting positive selection for outcrossing. But why did *M* sometimes fail to fix when outcrossing was initially extremely advantageous? Certainly, *M* was sometimes lost by chance before it had a chance to spread (Fig. 5). However, drift cannot explain why *M* was often rapidly lost after reaching high frequencies (red lines in Fig. 4, middle and right columns). What process determined whether invasion proceeded to fixation or was reversed?

For the outcrossing modifier to fail to fix on average after initially invading, either the expected fitness of selfed offspring must rise or the expected fitness of outcrossed offspring must fall. Several causes are possible, but not all were operating in our model. Outbreeding depression, in which outcrossed offspring are less fit than selfed offspring on average, and recombination load, in which recombination reduces fitness by breaking up fit haplotypes, are often caused by local adaptation and epistasis, respectively. We modeled neither local adaptation nor epistasis, however.

Three other processes were probably responsible for the eventual drop in the frequency of *M*. First, the difference between the expected fitness of outcrossed and selfed offspring fell as soon as *F*_1_s were generated—further outcrossing was less advantageous because it still draws from a pollen pool in which deleterious recessives shared with the *F*_1_s are common. Rapid reduction in the advantage of outcrossing probably explains why simulations with the smallest *s* did not result in fixation of *M*. Second, recombination in outcrossed individuals generated lightly loaded haplotypes that suffered less from selfing. These haplotypes were introduced into the selfing subset of the population because *Mm* individuals produced some *mm* seeds when they received *m* pollen. Third, selection among these recombinant haplotypes resulted in purging of deleterious alleles and an increase in the expected fitness of selfed offspring. That is, selfing generated linkage disequilibrium between the selfing allele and high fitness homozygous genotypes, in a manner similar to the way in which the “reduction principle” (Feldman and Liberman 1986) favors alleles that decrease the recombination rate because they generate LD with high fitness haplotypes.

We hypothesize that *M* went extinct unless it reached fixation before recombination and purging could produce sufficiently fit selfing genotypes. Figure 5 shows that purging was usually incomplete when *M* fixed but often complete when *M* was lost. This is consistent with purging favoring *m*. Purging was indeed most extensive in the longest simulations and least extensive in the shortest (Fig. S4). A race against time would also explain why simulations in which *M* fixed were often shorter than those in which m fixed (Fig. 5), and the sudden appearance of a sufficiently fit selfing genotype would explain why *M* abruptly declined from high frequency to extinction (Fig. 4). Additive outcrossing alleles fixed more quickly than dominant ones (Fig. 4) because selection continued to work against *m* even when it was rare and hence found mainly in heterozygotes. The increased fixation probability of additive outcrossing alleles was probably due to the reduced time spent segregating and thus the reduced opportunity for recombination to generate highly fit selfing genotypes (Fig. S3).

## Discussion

### Large heterosis favors outcrossing

Since heterosis can only be reaped through outcrossing, it provides an initial advantage to alleles that increase the rate of outcrossing in otherwise highly selfing populations. However, we found that this advantage was usually insufficient to drive such alleles to fixation. Across all of parameter space that we examined, the smallest amount of heterosis that resulted in the adaptive fixation of the outcrossing allele required the fitness of interpopulation heterozygotes to be 29.12 times that of individuals in either population. This results in 2812% heterosis, a level almost fortyfold larger than the highest heterosis we encountered in the literature (73.6% heterosis in a self-compatible population of *Leavenworthia alabamica*; Busch 2006) and over sixtyfold larger than the 42.6% heterosis for total survival documented in *Scabiosa columbaria* (van Treuren et al. 1993). The extent of heterosis required for the adaptive fixation of an outcrossing allele not only surpasses levels observed to date, but also likely exceeds any theoretically reasonable level.

### Strong selection required

Even a disadvantage per fixed deleterious allele of *s* = 0.1, the smallest we considered, is large enough that its fixation would be effectively impossible in nature. And yet at least thirty-two such fixations were required to have occurred in each population in order for the outcrossing mutation to fix more often than a neutral allele. The population-specific fixation of beneficial alleles with a similar magnitude is more plausible, but it is hard to believe that so many universally advantageous mutations that were possible in both populations would have only arisen in one. However, we have shown that it is at least theoretically possible for a modifier to invade populations subject to heterosis. Heterosis alone can be sufficient to favor the fixation of outcrossing modifier alleles even if purging has completely removed inbreeding depression.

Surprisingly, we find that an allele promoting outcrossing fixes more often at intermediate selection coefficients than at larger ones (Figs. 2a and 2b). How could increasing ***s*** (and thus the initial advantage of outcrossing) decrease the fixation probability of the outcrossing allele? Whitlock et al. (2000) found that intermediate selection coefficients maximize heterosis, but this is because they explicitly modeled the process by which heterosis accumulates. Decreasing *s* simultaneously allowed deleterious alleles to drift to high frequencies and reduces the heterosis contributed by those alleles in their model. The balance between these effects cannot explain our results, however, because we assumed that deleterious alleles began at high frequency regardless of their selection coefficients. Moreover, Whitlock et al. (2000) referred to intermediate selection coefficients that were two orders of magnitude smaller than the coefficients that are intermediate among our parameters. One possibility is that, although increasing *s* increased the initial magnitude of the load, it also hastened its purging. Intermediate *s* values might have balanced the magnitude and duration of the load in a way that maximized the advantage of outcrossing over the simulated period.

### Conditions favoring outcrossing in nature

Only unrealistically great heterosis was sufficient to favor outcrossing in our model, despite the fact that many of our assumptions were already very favorable to outcrossing. We assumed that the parent populations were of equal fitness (thus prolonging coexistence of the two initial genotypes), that inferior alleles were completely recessive (thus enlarging heterosis), and that there was no reproductive assurance (thus reducing the advantage of selfing). But did we neglect any plausible conditions that might favor outcrossing in nature?

#### Polygenicity of the load

The genetic basis of the load, not just its magnitude, determined the outcome of mating system evolution: polygenic loads favored outcrossing more than oligogenic loads of equal magnitude (Fig. 3). A possible explanation is that, if it is harder to purge many alleles than few, unpurged segregating deleterious alleles will generate inbreeding depression and favor outcrossing in the polygenic case. Simulating under loads of many more slightly deleterious alleles might be both more realistic and more likely to allow evolution of outcrossing, though some simplifications would need to be made because our current simulation scales poorly with number of loci. A large component of inbreeding depression in some outcrossing populations does indeed consist of many weakly deleterious alleles (Willis 1999), which are relatively unlikely to be purged during transitions to selfing.

#### Asymmetrical genotype frequencies

Nature differs from our main model in that there is no reason to suspect that the parental populations would be of equal size or that the initial genotypes would be at equal frequencies. When we varied the frequency of the two initial genotypes, we found that the outcrossing modifier fixed more often when it arose on the rarer genetic background (Fig. 6). This is likely a result of increasing the initial advantage of outcrossing: if the outcrossing individual seldom mates with similar genotypes at random because they are rare, the expected fitness of its collective offspring is greater. Unequal genotype frequencies could represent an outcrossing mutation arising after the two initial genotypes had drifted in frequency for several generations. Alternatively, the rarer outcrossing genotype could be an individual that arrived recently through a rare migration event.

The latter possibility is relevant to the putative reversions to outcrossing in the Hawaiian flora hypothesized by Baker (1967). We can construct a verbal model for the evolution of outcrossing on distant islands. Suppose that a single self-compatible individual has dispersed to a distant island and has established a population through selfing (Baker 1955). The combined effects of selfing and the demographic bottleneck have exposed deleterious recessive alleles, which were then purged. However, the population has also fixed a local drift load. A highly outcrossing conspecific migrant then arrives from the mainland. If this migrant had arrived alone, it would lack any mates and would produce no offspring. But since it arrived after a predominantly selfing population had already taken hold, these selfing individuals could act as potential mates. Since a selfing individual only benefits from the transmission advantage of selfing if it also continues to act as an outcross sire, it is conceivable that the selfing residents would retain the ability to fertilize the outcrossing migrant. (Of course, if high levels of selfing in the population substantially reduced the probability of successful outcross siring, selection would have favored reduced investment in pollen export. This scenario requires either that the island residents are partially but not fully selfing, or that outcrossing migrants arrive before selection can abolish the ability of the residents to sire outcross offspring.) Hybridization between the outcrossing migrant and the predominantly selfing residents will simultaneously introduce alleles that increase the outcrossing rate and superior alleles that can mask deleterious recessives. Selection may then favor outcrossing mechanisms, possibly including dioecy.

However, other historical scenarios could also explain the prevalence of dioecy on islands. First, dioecy was likely the ancestral state for some taxa before dispersal to an island (Sakai et al. 1995). Even in cases in which it evolved *in situ*, dioecy could have evolved for reasons unrelated to inbreeding avoidance (Charnov 1979; Bawa 1980; Givnish 1982). Islands could also have been colonized without purging either because a substantial component of inbreeding depression was weakly deleterious and resistant to purging (Willis 1999; Keller and Waller 2002), or because the initial population was large enough that many deleterious alleles remained masked in heterozygotes (e.g., many individuals arrived on rafts or in many-seeded fruits).

#### Inbreeding depression in ancestral populations

The ancestral populations in our model started with no within-population inbreeding depression because they were completely monomorphic. In nature, however, moderately and even highly selfing populations show inbreeding depression (Winn et al. 2011), which may sometimes be large (Herlihy and Eckert 2002, 2004). This implies either that these selfing populations have not fully purged their inbreeding depression or else that they have accumulated inbreeding depression. If alleles underlying inbreeding depression are individually weakly deleterious, then more of them will persist even after being exposed to selection by selfing (Charlesworth et al. 1990). An outcrossing population of *Mimulus guttatus* retained most of its inbreeding depression for many characters after five generations of artificially enforced selfing, which suggests that weakly deleterious alleles make up a large fraction of inbreeding depression in this population (Willis 1999). Inferior recessive alleles at loci in repulsion-phase linkage disequilibrium (pseudo-overdominance) also resist purging because the superior double homozygote cannot be generated until the linkage disequilibrium is broken by recombination (Charlesworth and Willis 2009). Heterosis and remaining inbreeding depression could together favor outcrossing under a broader array of conditions. If so, evolving greater outcrossing might not require as high levels of heterosis as our model suggests.

#### Pollen discounting

We assumed pollen discounting was absent, but this may be a crucial omission. Pollen discounting is the degree to which a trait that increases self-fertilization also reduces outcross siring success. When pollen discounting is absent, selfing transmits 50% more gametes than outcrossing (Fisher 1941) because seeds sired through selfing are in addition to those sired through outcrossing. Increasing pollen discounting reduces the transmission advantage, and complete pollen discounting totally eliminates it (Nagylaki 1976). By reducing the advantage of selfing, pollen discounting could greatly expand the parameter space in which the outcrossing allele is favored. Although we can qualitatively predict that increasing pollen discounting will increase the outcrossing allele’s fixation probability, we do not know the rate of this increase. Since purging was able to eliminate even an initial disadvantage of selfing far exceeding the transmission advantage in our simulations, we expect the outcome to be insensitive to all but high levels of pollen discounting.

### Maintenance of heterosis

Heterosis was rarely sufficient to allow the evolution of outcrossing because it promoted its own erosion through introgression. If heterosis has actually promoted the evolution of outcrossing from predominant selfing in nature, then some other factor likely prolonged its existence. Bierne et al. (2002) showed that genetic barriers to hybridization could reduce introgression and prevent the homogenization of populations that eliminates heterosis. Partial physical (e.g., distance) or biological (e.g., prezygotic isolation) barriers could also maintain heterosis. All these barriers would decrease the magnitude of the advantage of outcrossing by making hybrids rarer or less fit, but they would also extend the advantage’s duration by maintaining the allele frequency differences underlying heterosis. Our results showed that even enormous initial heterosis was rarely sufficient for outcrossing to evolve because it decayed quickly. A small but lasting heterosis maintained by genetic or physical barriers might more realistically allow outcrossing to evolve than did the large but transient heterosis of accumulated fixed differences we modeled.

### Adaptive introgression

Although we focused on the evolution of outcrossing, we also found that secondary contact often greatly increased mean fitness even in cases in which the outcrossing modifier was ultimately lost (Fig. 5). Hybridization allowed the adaptive introgression of superior alleles from each parental population. Recombination was apparently sufficient to eliminate the initial association between superior and inferior alleles more quickly than selection could produce linkage disequilibrium. Sorting out superior alleles from inferior ones may be essential to adaptive introgression in nature. For example, positive selection on sequences introgressed from Neanderthals into humans seems to have occurred at some loci despite apparent selection against Neanderthal ancestry across most gene-rich regions of the genome (Sankararaman et al. 2014). An allele that is inferior in the recipient population may have persisted in the donor population for several reasons, such as local adaptation to the donor’s environment or small donor effective population size. Juric et al. (2016) and Harris and Nielsen (2016) hypothesized that the deleterious alleles were effectively neutral in the small populations of Neanderthals, but became exposed to selection in the larger human populations. Whatever the source of the deleterious alleles, advantageous alleles were able to escape this genetic background through recombination. In our model, the lowered effective recombination rate in highly selfing populations was not enough to prevent a similar process. Introgression may therefore be an important source of new advantageous alleles in predominantly selfing populations, but only if they occasionally outcross. This occasional outcrossing might be due to transient invasions of outcrossing alleles, as in our model, or it might be a result of a stable partially outcrossing mating system.

### Alternative pathways for reversion to outcrossing

We proposed heterosis as an alternative advantage of outcrossing that could exist even in purged, selfing populations. However, other, potentially less transient advantages also exist. The advantage of recombination (and thus of outcrossing) could increase when the population encounters a new, co-evolving natural enemy such as a pathogen. Rare recombinant genotypes to which the pathogen has not adapted might have exceptional fitness (Levin 1975; Jaenike 1978; Bell and Smith 1987; but see Otto and Nuismer 2004). Such an advantage could conceivably persist indefinitely as long as the host and pathogen populations coexist. This situation might correspond to the introduction of a biocontrol agent to an invasive selfing weed that previously lacked local natural enemies. Another possibility is for an environmental shift to increase inbreeding depression. Inbreeding depression is known to be dependent on the environment (Keller and Waller 2002). For example, inbreeding depression might manifest in a plant’s natural habitat but not in the artificially benign environment of a greenhouse. Because purging only removes alleles that were deleterious in the environment in which purging occurred, an environmental shift can cause unpurged conditionally neutral alleles to become deleterious (Bijlsma et al. 1999) and potentially contribute to inbreeding depression. If shifts to new environments were to occur often enough, repeatedly regenerated inbreeding depression could favor the evolution of outcrossing. Yet another possibility for an increased net advantage of outcrossing is a reduction in the advantage of selfing. For example, animal pollinators that had previously declined in a population may rebound, thus reducing the reproductive assurance provided by selfing. This last scenario would need to be paired with another advantage of outcrossing because the transmission advantage of selfing would remain even if reproductive assurance were to fall to zero (as it does in our model).

### Conclusion

The reversion from predominant selfing to outcrossing remains a theoretically neglected process partly because predominantly selfing populations lack much of the allelic diversity that normally favors outcrossing. Heterosis from local drift load circumvents this barrier because within-population variation is regenerated from among-population divergence. However, the architecture of local drift load is unlikely to maintain an advantage of outcrossing for long. Nevertheless, secondary contact may promote adaptive introgression even if greater levels of outcrossing do not evolve. Future investigations should search for advantages of outcrossing that both survive purging and that persist long enough for outcrossing alleles to invade.

## Acknowledgments

We would like to thank the labs of Y.B., E.E.G., and Suzanne McGaugh for their critique of this manuscript. A.H. was funded by the Graduate Excellence Fellowship from the Department of Ecology, Evolution, and Behavior, University of Minnesota during this work.

## Appendix Appendix: Condition for increase of the outcrossing modifier allele with two viability loci

Here, we derive the values of the selection coefficient, *s*, for which the frequency of the outcrossing modifier, *M*, initially increases in frequency. We assume here two viability loci, which, when combined with the three mating system genotypes, yields 27 genotypes to track.

The frequency *p* of the outcrossing allele increases to *p*′ in the next generation when

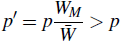

where *W*_*M*_ is the marginal fitness of the outcrossing allele and 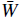 is the mean fitness in the population. Multiplying both sides by 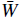 condition becomes and expanding *pW*_*M*_ into a weighted average of genotype frequencies, this condition becomes

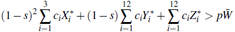

where *s* is the selection coefficient for each viability locus, the *X*_*i*_ are the frequencies of the three genotypes homozygous for the inferior allele at both viability loci, the *Y*_*i*_ are the frequencies of the twelve genotypes homozygous for a single deleterious allele, the *Z*_*i*_ are the frequencies of the twelve genotypes homozygous for the deleterious allele at neither viability locus, *c*_*i*_ is the proportion of *M* alleles (0, 0.5, or 1) at the modifier locus in the *i*th genotype, and * denotes a frequency after mating but before selection. Expanding

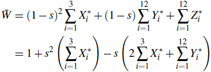

and substituting

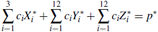

and collecting terms in powers of *s*,

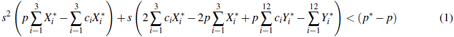

Given all genotype frequencies, this quadratic inequality can be solved to determine the values of *s* for which *p′ > p*. When there are no double homozygotes for the inferior alleles, as is the case before *F*_2_s and backcrosses are generated, the *s*^2^ term becomes zero, the inequality becomes linear, and the invasion condition is

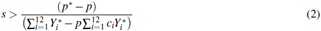

**Figure S1:**
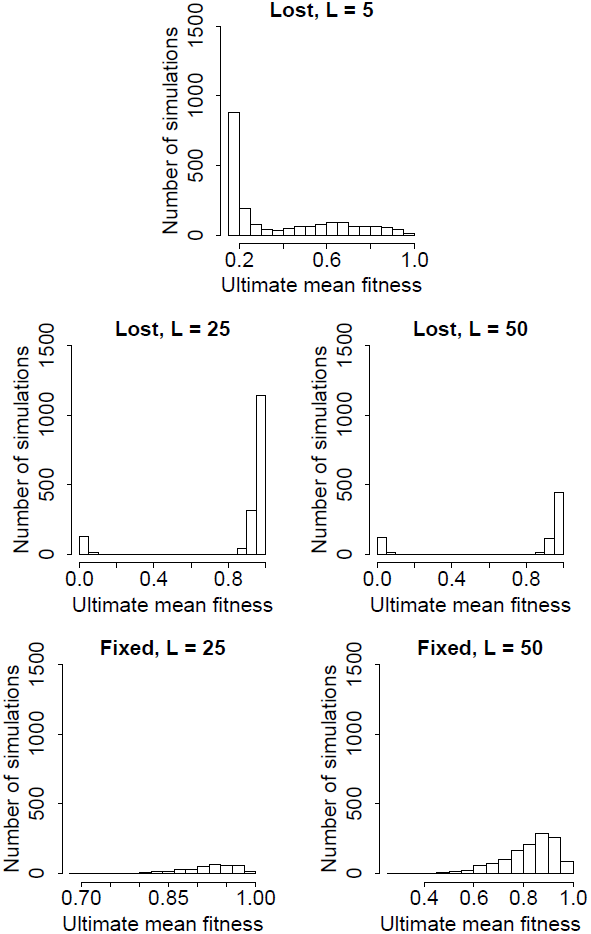
Distribution of final mean population fitness at the end of the simulation. Fitness was not tracked for the simulations in which *M* was additive, so *M* is dominant for all parameter combinations. *M* never fixed for *L* = 5. For all parameter combinations, *s* = 0.3. The parameter combinations in which *L* = 50 are identical to the main panels of Fig. 5. “Lost” and “fixed” in panel titles refer to the fate of *M*.

**Figure S2:**
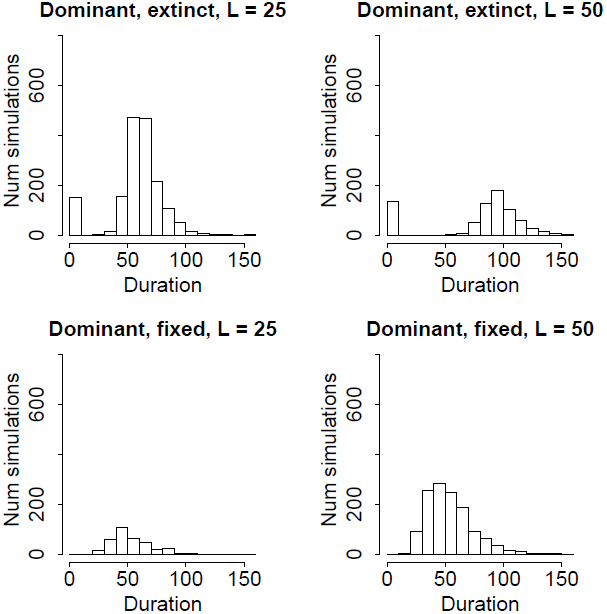
Distribution of simulation durations (dominant outcrossing allele). For all parameter combinations, *s* = 0.3. The parameter combinations in which *L* = 50 are identical to the insets of Fig. 5. “Extinct” and “fixed” in panel titles refer to the fate of *M*.

**Figure S3:**
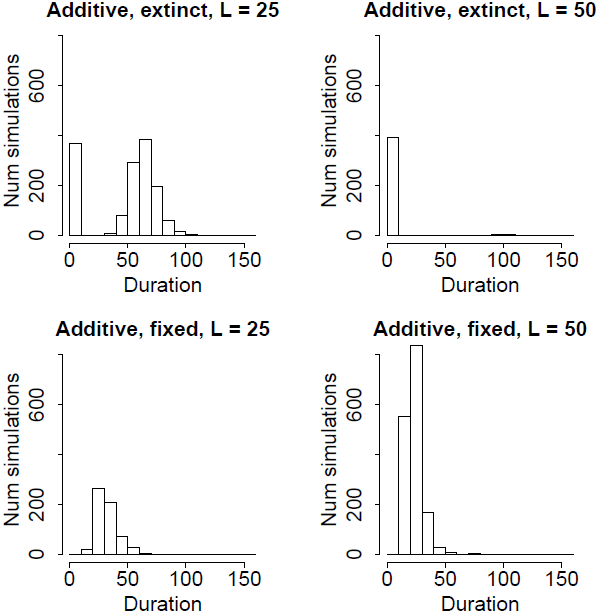
Distribution of simulation durations (additive outcrossing allele). For all parameter combinations,*s* = 0.3. “Extinct” and “fixed” in panel titles refer to the fate of *M*.

**Figure S4:**
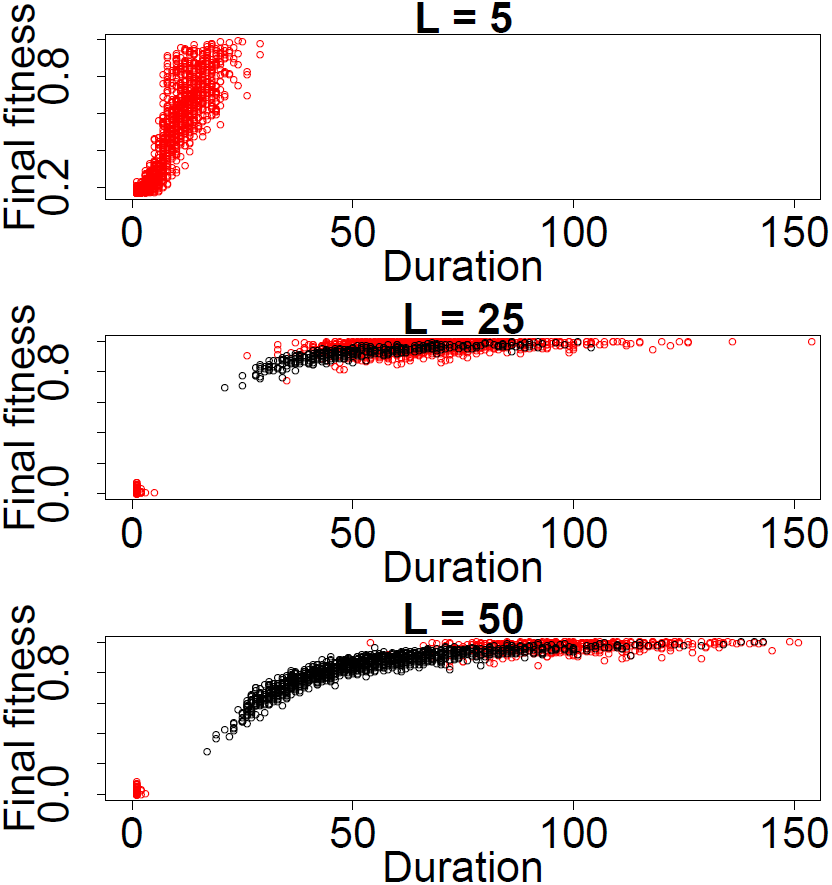
Extent of purging. The mean fitness of the population at the end of the simulation is plotted against the duration of the simulation. Black circles represent simulations in which *M* fixed, and red circles represent simulations in which *M* was lost. Purging was more extensive in the simulations that lasted longer. Very little purging occurred in the shortest simulations. *M* tended to fix in the simulations of intermediate duration and incomplete purging. Parameter combinations with different *L* are plotted separately. Data are identical to those of Fig. S1 and Fig. S2. *M* is dominant and *s* = 0.3 for all parameter combinations.

